# Implementation of a pre- and in-scan system to minimize head motion in pediatric participants undergoing fMRI scans

**DOI:** 10.1101/2020.03.04.975417

**Authors:** Corey Horien, Scuddy Fontenelle, Kohrissa Joseph, Nicole Powell, Chaela Nutor, Diogo Fortes, Maureen Butler, Kelly Powell, Deanna Macris, Kangjoo Lee, James C. McPartland, Fred R. Volkmar, Dustin Scheinost, Katarzyna Chawarska, R. Todd Constable

**Affiliations:** Interdepartmental Neuroscience Program, Yale School of Medicine, New Haven, CT, USA; Yale Child Study Center, New Haven, CT, USA; Department of Radiology and Biomedical Imaging, Yale School of Medicine, New Haven, CT, USA; Department of Psychology, Yale University, New Haven, CT, USA; Department of Statistics and Data Science, Yale University, New Haven, CT, USA; Department of Pediatrics, Yale School of Medicine, New Haven, CT, USA; Department of Neurosurgery, Yale School of Medicine, New Haven, CT, USA

**Keywords:** head-motion, mock scan, pediatric imaging, functional connectivity, autism spectrum disorder

## Abstract

**Background:** Performing fMRI scans of children can be a difficult task, as participants tend to move while being scanned. Head motion represents a significant confound in functional magnetic resonance imaging (fMRI) connectivity analyses, and methods to limit the impact of movement on data quality are needed. One approach has been to use shorter MRI protocols, though this potentially reduces the reliability of the results.

**Objective:** Here we describe steps we have taken to limit head motion in an ongoing fMRI study of children undergoing a 60 minute MRI scan protocol. Specifically, we have used a mock scan protocol that trains participants to lie still while being scanned. We provide a detailed protocol and describe other in-scanner measures we have implemented, including an incentive system and the use of a weighted blanket.

**Materials and methods:** Participants who received a formal mock scan (n = 12) were compared to participants who had an informal mock scan (n = 7). A replication group of participants (n = 16), including five with autism spectrum disorder, who received a formal mock scan were also compared to the informal mock scan group. The primary measure of interest was the mean frame-to-frame displacement across eight functional runs during the fMRI protocol.

**Results:** Participants in the formal mock scan and replication group tended to exhibit more low-motion functional scans than the informal mock scan group (*P* < 0.05). Across different functional scan conditions (i.e. while watching movie clips, performing an attention task, and during resting-state scans), effect sizes tended to be large (Hedge’s *g* > 0.8).

**Conclusion:** Results indicate that with appropriate measures, it is possible to achieve low-motion fMRI data in younger participants undergoing a long scan protocol.

## Introduction

Functional magnetic resonance imaging (fMRI) has proven to be a powerful tool to study brain function. A promising approach using fMRI data is to measure functional connectivity, in which time courses of the blood oxygen level-dependent (BOLD) signal are correlated across regions of interest [1], and to use this connectivity data to develop functional phenotypes. Such analyses have been used extensively to characterize the brains of younger children [2–6] including those with autism spectrum disorder [6–10], and the hope is that one day such functional connectivity approaches can be used to help individual patients in clinical settings [11–13].

Nevertheless, scanning children can be challenging, especially because younger children tend to move while being scanned. The effect of motion on measures of functional connectivity is well documented [14, 15], and it can introduce major confounds into analysis pipelines. While there are post-hoc methods to clean fMRI data to reduce the impact of motion [16–21], there is no consensus about the best way to do so [22, 23].

An additional and complimentary approach is to decrease in-scanner motion in this population during data acquisition. A common strategy with children is to use shortened scan protocols and prioritize the scans of interest—for example, collecting only a structural scan for common-space template registration and only 1-2 resting state or task-based scans. However, numerous groups have shown that the reliability of functional connectivity measures increases with increasing scan duration [24–28], thus choosing a shortened scan protocol increases the risk of obtaining unreliable results.

In addition, other in-scanner methods exist to help obtain low motion data. For instance, one approach, Framewise Integrated Real-time MRI Monitoring (FIRMM), analyzes head motion in real-time and allows the scanner operator to collect data until a satisfactory amount of low-motion data have been obtained [29]. This method is being utilized by investigators at certain sites in the Adolescent Brain Cognitive Development (ABCD) study [30]. Other groups have demonstrated that showing movies—actual movie clips with and without feedback [31] and low-demand clips of abstract shapes (e.g., *Inscapes*; [32])—during a scan can reduce motion in younger children. While these approaches have proven useful, there are potential issues with each. For instance, it would be difficult to use FIRMM when completing task-based scans, as differences in task length might affect task performance and confound analyses. Showing participants movie clips affects connectivity, even when using a low-demand movie such as *Inscapes* with minimal semantic content [31, 32]. Because of the impact on connectivity, it would be preferable to decrease motion without introducing confounds.

Mock scanning protocols are one potential solution to the problem of in-scanner motion. While exact details vary, this approach typically entails placing participants in an environment designed to mimic the real scanning environment, desensitizing them to the scanning experience, and training them to limit movement. Numerous groups have described successful implementation of a mock scan protocol and have shown that it can be used to limit in-scanner motion in younger children [33–37]. However, these studies have tended to be shorter in duration (i.e. between 20-45 minutes to scan one participant) and have tended to collect only structural and/or a few task or resting-state scans, so the efficacy of mock scanning when using longer MRI protocols is unclear. In addition, most of these studies have lacked a control group (e.g., a group that was part of the same study but did not receive a mock scan), rendering effectiveness of the mock scan unquantifiable.

Here we describe methods our group has used to decrease in-scanner motion in pediatric participants undergoing a longer MRI protocol (61 minutes total scan time; 80-90 minutes with eye-tracker set-up, time between scans, etc.). We detail the steps of a mock scan protocol we have developed and describe other in-scanner steps to limit motion, including an in-scan incentive system and use of a weighted blanket. We show that these steps significantly reduce head motion during functional scans compared to a group that did not receive a formal mock scan, did not use the weighted blanket, and did not participate in the in-scan incentive system. Further, we have validated our findings by training a separate group of researchers to conduct the mock scan protocol for a separate group of participants, as well as interact with the child during the actual fMRI scan protocol. We found that motion was again low in the functional scans in the replication group of participants, including some with autism spectrum disorder (ASD). Our data suggest that by using a formalized mock scan protocol, in conjunction with other in-scan steps, investigators can achieve low motion scans in pediatric participants undergoing a long fMRI protocol.

## Materials and methods

### Participants

This study was approved by the Yale Institutional Review Board. Informed consent was obtained from the parents of participants. Written assent was obtained from children ages 13 – 17; verbal assent was obtained for participants under the age of 13. As appropriate, we have adhered to the Strengthening the Reporting of Observational Studies in Epidemiology (STROBE) guidelines for reporting results related to observational studies [38].

This sample was derived from an ongoing study being conducted to test brain-wide, network-based functional connectivity models of ASD. Participants were screened over the phone for basic developmental history and MRI safety factors. Children and adolescents with history of prematurity, known genetic abnormalities, or an IQ below 70 were not invited to participate in the study. Participation included two visits, the first for clinical assessments and the second for fMRI scanning. On the day of the clinic visit the participants underwent diagnostic characterization examining severity of developmental abilities using the Differential Ability Scales-II (DAS-II [39]) and autism symptoms using the Autism Diagnostic Observation Schedule-2 (ADOS-2 [40]). Diagnostic classification was based on clinical best estimate diagnosis by a team of clinical psychologists based on test results and any available reports of developmental and medical history (see Table 1 for demographic information).

**Table 1.** Demographic information.

| **Measure: group means (s.d.)** | **Informal mock scan group** | **Formal mock scan group** | **Replication group** | ***P*-value (formal mock scan group vs. informal mock scan group)** | ***P*-value (replication group vs. informal mock scan group)** |
| --- | --- | --- | --- | --- | --- |
| Number of participants | 7 | 12 | 16 | - | - |
| Number of participants with ASD | 0 | 0 | 5 | - | - |
| Age in years | 10.24 (2.05) | 11.31 (2.78) | 12.14 (2.89) | 0.39 | 0.13 |
| Males per group | 6 | 6 | 5 | 0.32 | 0.03 |
| IQ: verbal | 118.28 (10.81) | 117.5 (10.74) | 114.06 (12.22) | 0.88 | 0.44 |
| IQ: non-verbal | 116.43 (13.25) | 108.9 (10.72) | 108.81 (12.01) | 0.23 | 0.19 |
| IQ | 119 (15.44) | 113.08 (12.9) | 111.56 (11.25) | 0.38 | 0.21 |

Three groups are considered in the present study: a group who received a formal mock-scan, a group who received an informal mock scan, and a group who received a formal mock scan led by other researchers (this group served as the replication group; details below). Participants were assigned into each group based on when they enrolled in the study. That is, the first seven participants received an informal mock scan, the next 14 participants received a formal mock scan, and the next 16 participants received a formal mock scan led by other staff members.

The first group of participants receiving a mock scan (n = 14; none with ASD) took part in a formal mock scan with intensive feedback. The corresponding author (CH) primarily led the mock scans with assistance from one of several research staff members (NP, KJ, CN, SF). In the mock scan group, the corresponding author was also responsible for leading the fMRI scans (e.g. communicating with the participant in between scans, task/eye-tracker setup, etc.), with the assistance of an MRI technologist, along with another research staff member (NP, KJ, CN, SF). We note that for training purposes the final three mock scans were led by either NP, KJ, or CN, while CH continued to lead the actual MRI session, setup, and in-scan incentive system for these participants. We note that two out of the 14 (14%) of the participants in this group were unable to complete the mock scan due to discomfort. These did not take part in the actual MRI session and were excluded from the study; hence, the final sample size of the mock scan group was n = 12.

The group of participants receiving an informal mock scan (n = 7; none with ASD) took part in the following procedure: After a 2-3 minute period of desensitization (i.e. laying on the mock scan table outside of the mock bore), participants simply lay in the mock scanner for 5-10 minutes while gradient sounds from the scanner were played (the exact time depended on their comfort level—if participants said they felt comfortable after 5 minutes, the session was concluded). Participants were given feedback for gross head movements or other large movements. No other specific feedback was given. The corresponding author (CH) conducted these scans, along with SF and KJ. The informal mock scan group did not use a weighted blanket during the scan, and participants did not partake in the in-scan incentive system. No participant dropped out before or during the scan; hence, the final sample size was n = 7 for this group.

The second group of participants receiving a formal mock scan (n = 16; 5 with ASD) took part in a mock scan with intensive feedback led by two other members of the research staff (NP, KJ, CN, DF, MB); two members of the research staff were also responsible for running the fMRI scans with the assistance of an MRI tech. The purpose of this group was two-fold: first, to determine if low-motion data could be obtained in a group of participants who had not been primarily trained by CH, and second, to determine if the protocol could be used in individuals with ASD.

Both groups receiving a formal mock scan also used a weighted blanket during the scan and participated in an in-scan incentive system (which we detail below in the “Additional steps to limit motion” section in the Methods). For concision, we refer to these groups hereafter as the “formal mock scan” and “replication” groups, though use of these other factors to limit motion must be kept in mind. We hereafter refer to the group that received the informal mock scan as the “informal mock scan” group.

We note that when comparing these groups, we are not interested in the individual effect of each step on motion (i.e. whether the mock scan protocol, the weighted blanket, or the in-scan incentive system has a bigger effect relative to the other). Rather, we are simply interested if all three factors result in lower motion functional data in the formal mock scan and replication groups compared to the informal mock scan group.

### Description of mock-scanning environment

The mock scanner consisted of a 5-foot-long motorized bed, a wooden mock head coil, and an audiovisual presentation system (Fig. 1). Speakers placed in corners of the room were used to play gradient sounds. To mimic the scanning environment as closely as possible, participants were given earplugs to wear; auditory stimuli were delivered through headphones. A fan inside the bore was also turned on to replicate the scanning environment. Visual stimuli were seen through the use of a mirror system that displayed visual stimuli situated at the back of the bore. MoTrak (https://pstnet.com/products/motrak/) was used as the head motion tracking system in this study (Fig. 2), a system designed to allow participants to see how their head moves in real-time while they are in the mock scanner. Head movement was monitored through the use of a MoTrak headband. In general, two staff members conducted a mock-scan: one interacted with the child and provided feedback; the other staff member set-up and monitored the MoTrak software. The staff member monitoring the MoTrak software was able to view the participant through a camera to ensure participant compliance and observe how the participant responded to feedback.

**Fig. 1.**
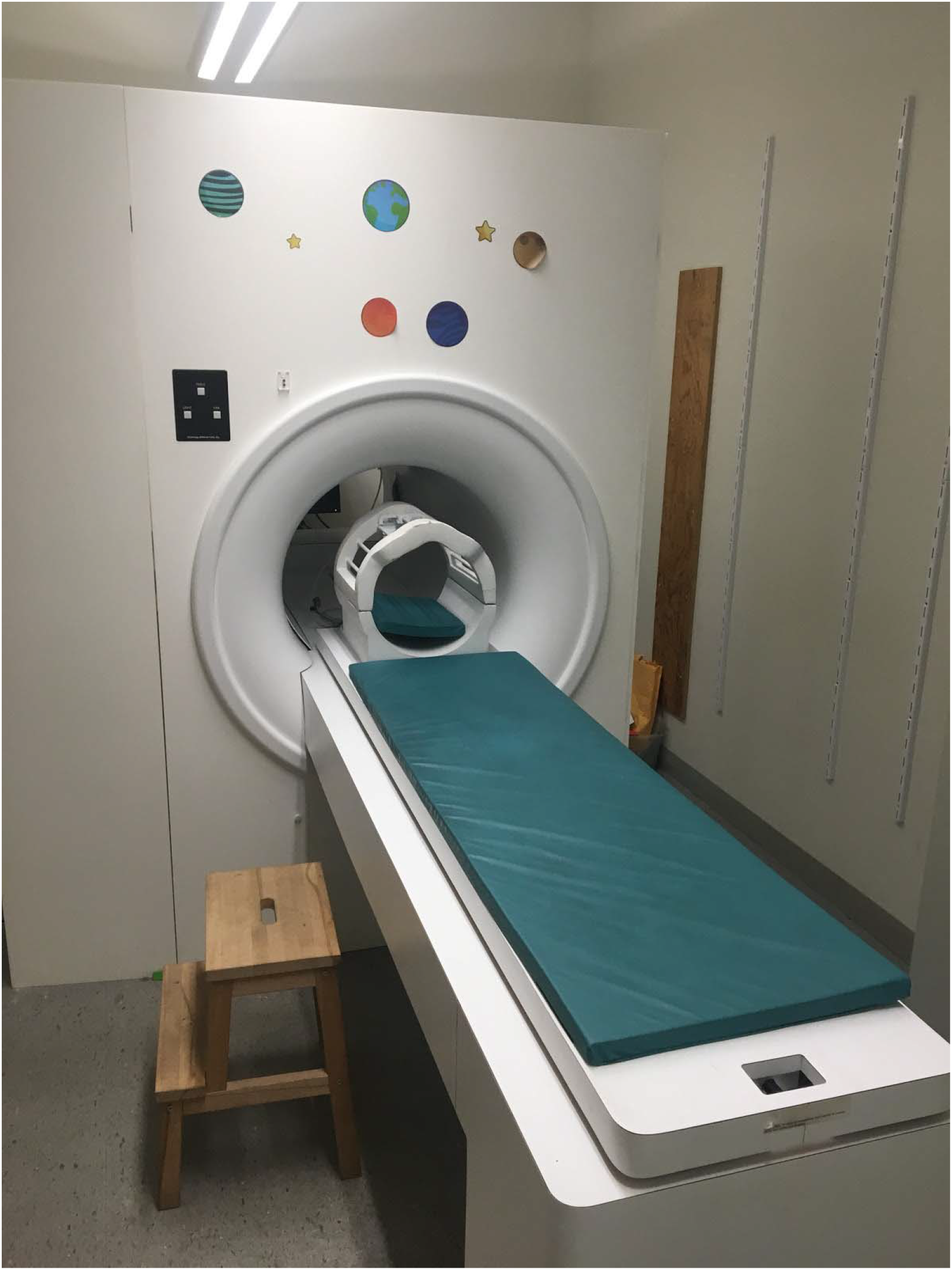
The mock-scanner used in this study. A motorized bed is used to move the participant back into the bore. The participant views visual stimuli through the use of a mirror on the head-coil. The participant wears headphones to hear sounds from the movie. Speakers (not visible in the photo) are used to play gradient sounds while the participant lay in the mock-scanner

**Fig. 2.**
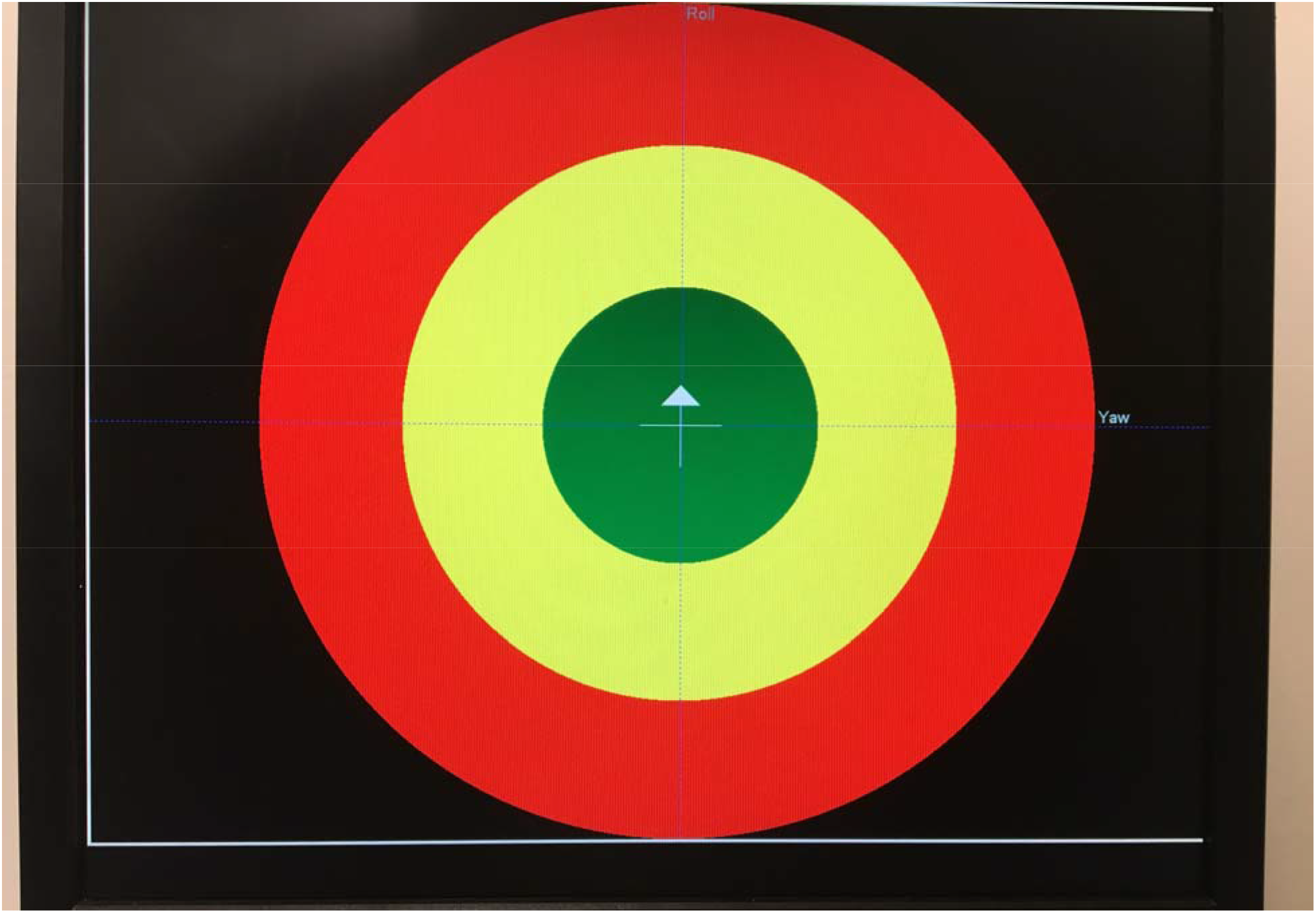
The target system used in conjunction with MoTrak software. The cursor (shown in the center of the target as a cross) tracks participant head movement. This screen is visible to the participant at the back of the mock scanner through a mirror system. Head movement results in the cursor moving on the screen—hence, the participant can see how their movement causes the cursor to move. Participants are encouraged to stay in the green circle as much as possible. Depending on the step of the mock scan protocol (see below for more details), verbal feedback is also given to the participant (“You moved your arms, and it caused your head to move; did you see that on the screen?”)

**[Figure removed per bioRxiv confidentiality guidelines]**

**Fig. 3.** Single frame from the direct gaze with speech condition of the selective social attention task. Four conditions were used: direct gaze with speech, direct gaze with no speech, no direct gaze with speech, and no direct gaze with no speech

### Description of formal mock scan protocol

The mock scan protocol consists of nine steps, including several mock scan runs with incremental incorporation of visual stimuli, scanner sounds, and feedback about motion. The entire sequence takes about 45-50 minutes to complete, and was typically conducted four days before the actual MRI protocol, depending on participant availability.

The first step of the mock scan involved introducing the participant to the mock scan environment and was three minutes long (Table 2). The participant was allowed to walk around the room, look inside the mock scanner, and ask any questions. Age-appropriate language was used to describe the mock scanner (e.g., for younger participants: “This is a camera we use to take pictures of your brain.”). The goals of the mock scanning session were explained to the participant. (“It is important that when you are in the scanner, you stay very still. We are going to practice that over the next 45 minutes.”) As appropriate, a question and answer approach was utilized by the research staff with the participant: “What happens if you move around a lot when your picture is being taken? That’s right; it is blurry. That is why we need you to stay still when we are taking pictures of your brain.”

**Table 2.**
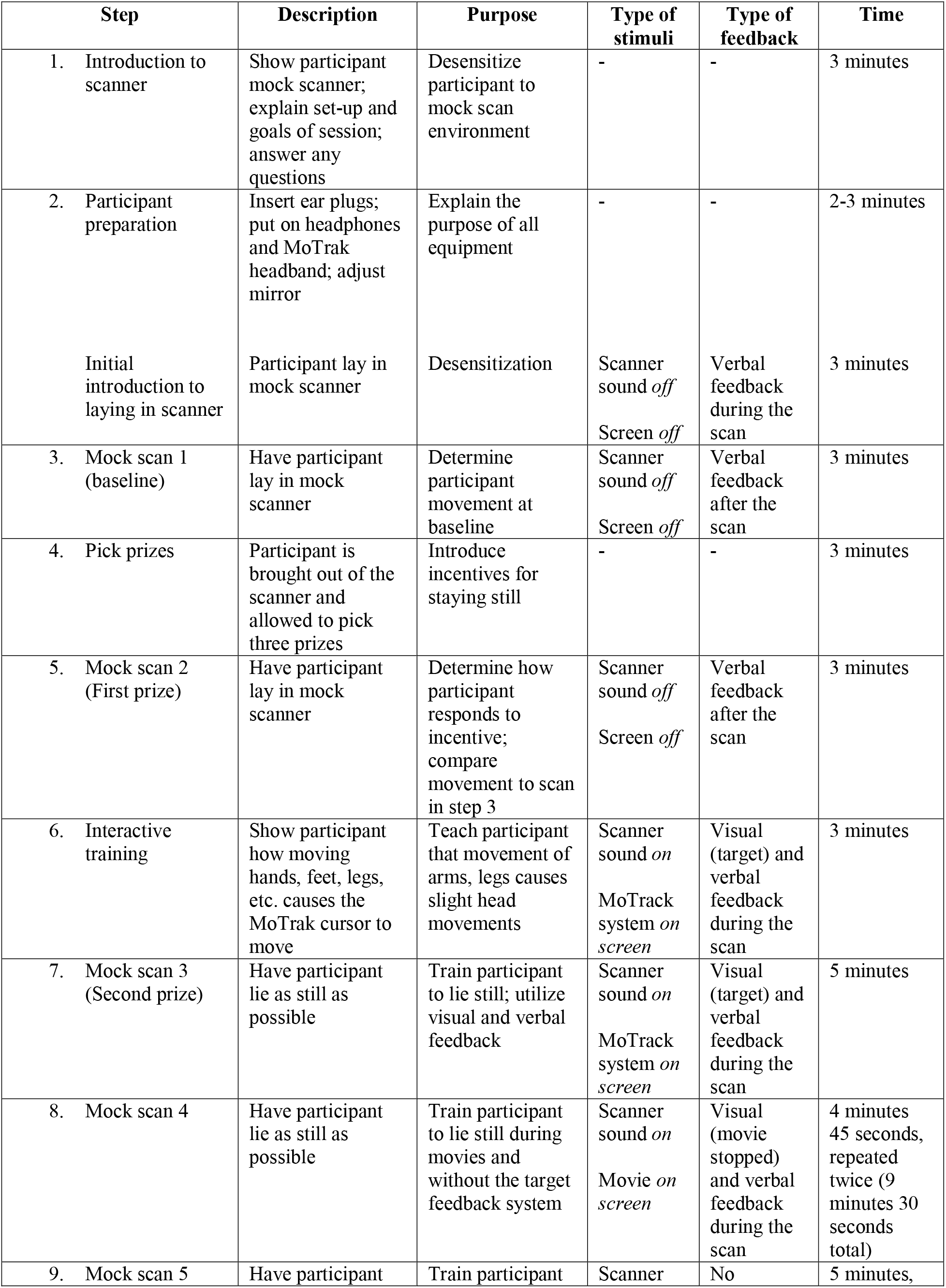
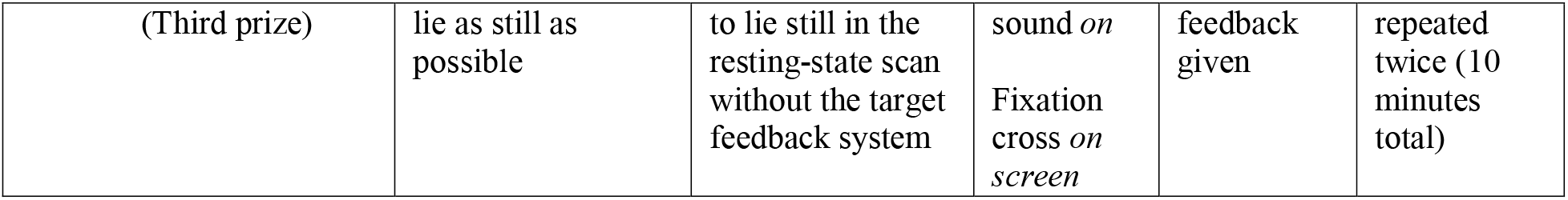
The steps used in the mock scan protocol.

The second step included preparing the participant for the mock scan. Ear plugs were given to the participant; after inserting them into the ears, headphones were applied over the ears by research staff, again with the intention of simulating the actual scanning environment as closely as possible. The MoTrak head-tracking headband was applied. The participant then laid back, and the mirror was adjusted so the participant could view the computer monitor placed at the back of the mock scanner. As appropriate, visual stimuli (including the target system from MoTrak, movie clips, or a fixation cross) were projected through the back computer monitor.

After this, the participant lay down in the bore of the mock scanner for three minutes. No visual stimuli were presented. No scanner sounds were used. The purpose of this step was to acclimate the participant to the feeling of lying down with the mock head coil in the scanner. Specific feedback based on direct observation of the participant was given by research staff for head and/or body movements. (“You’re moving your head right now—remember, our pictures are blurry if you do this.”) Positive feedback was given as appropriate. (“You’re doing such a good job staying still!”)

The third step consisted of testing how well the participant did in the mock scanner without feedback for three minutes. The purpose was to determine how much the participant moved at baseline without training. MoTrak software (at this point only visible to research staff) was used to record the amount of participant head-movement. No scanner sounds were used in this step. After three minutes, the participant was given feedback about motion throughout the scan. (“You did a good job keeping your arms still, but you moved your feet throughout the scan, and this caused your head to move. Remember, we want your entire body to stay still so we can take nice pictures of your brain.”)

In the fourth step, the participant was brought out of the scanner and an incentive was introduced. Specifically, participants were shown a collection of toys and were allowed to choose three. It was explained that if they stayed still throughout the scan, they would have a chance to win all three based on limiting motion during the rest of the procedure. The purpose of introducing the incentive was to determine how the participant responded to this motivating factor and to determine how this changed head motion compared to the baseline mock scan obtained during step three. For example, if the participant exhibited significant head movement during the mock scan in step three but then reduced head motion in response to the prizes, this was noted by the research staff, and the prize system during the actual MRI protocol (described below in the “Additional steps to limit motion” section) was adjusted accordingly, typically by allowing a participant to win two extra prizes per scanning session.

In the fifth step, the participant was put back into the mock scanner for three minutes, and head motion was tracked without sounds. If the participant moved their head fewer times than the mock scan in step three, they could win their first prize. No feedback was given by research staff during the scan; feedback about the total amount of head movement was given after the scan. (“You moved out of the green circle only two times that scan, so you win your first prize, but we noticed you still moved your feet quite a bit. Remember we want you to stay as still as a statue.”)

In the sixth step, research staff worked with the participant to show the effects of movement. For example, staff would have the participant look at the target and observe how much the cursor moved on the target in response to different movements of the participant’s hands and feet; after yawning; after taking a deep breath; after itching/fidgeting, etc. (“When you move your hands, your head also moves out of the green circle—did you see that? This is why we want your entire body to be still in the scan.”) The purpose of this step was to demonstrate what it means to lie still and to show that even though the participant might think they are laying still, they often might be moving in subtle ways that cause head movements.

In the seventh step, the participant completed a mock scan for five minutes with scanner sounds and with the MoTrak target system visible to the participant. Research staff gave specific verbal feedback to the participant throughout the scan, approximately every 20-30 seconds. (i.e. “You’re moving your hands a lot and this is causing your head to move out of the green circle. Remember we want your entire body to be as still as a statue.” “You’re doing such a good job lying still!”) The purpose of this step was to simulate a real scan through use of the scanner sounds and to train the participant to lie still. Before this mock scan, it was explained to the participant that if they stayed as still over the next three scans as they did during the fifth scan (i.e. moved out of the green circle fewer times) and if they responded to verbal feedback, they could win their second prize.

In the eighth step, the participant completed a mock scan for 4 minutes and 45 seconds while watching a movie clip with scanner sounds being played. This step was completed twice. The target system was no longer visible to the participant, though head movement was still tracked through the MoTrak system. The MoTrak system was configured so that every time the participant moved out of the green circle, the movie paused. Feedback was then given to the participant. (“Did you see that the movie stopped? This was because you were wiggling your hands and it caused your head to move. Remember to keep your hands still.”) Feedback was given between the first and second scan about the total number of times the participant moved out of the green circle.

In the ninth step, the participant completed a five-minute mock resting-state scan with scanner sounds. This step was completed twice. A fixation cross was shown to the participant in place of the MoTrak target. Head movement was still tracked through MoTrak. It was explained to the participant that no feedback would be given during the scan, that research staff would record the number of times the cursor moved out of the green circle, and that feedback would be given in between scans. It was also explained that the purpose of this step was to simulate what it would be like during a real scan (i.e. research staff would not be able to talk to the participant during the scan; there would be no target system presented during the actual scans; feedback would only be given in between scans, etc.) Finally, research staff explained that if the participant moved out of the green circle fewer times during each rest scan than they did in the second movie clip and if they responded to verbal feedback, they could win their third prize.

### Additional steps implemented during the MRI scan to limit motion

In addition to the mock scan protocol, we took other measures to limit motion in the mock scan group. We used an MRI compatible weighted blanket during the actual MRI session (https://www.mosaicweightedblankets.com/). We customized the blanket so it would be appropriately sized for children (blanket dimensions: 38” × 60”; eight pounds of total weight).

We also implemented a prize system during the actual MRI session with the mock scan group. Participants could win up to three prizes (toys, stickers, pencils, etc.) of their choice if they were deemed to have low motion scans. In general, a participant could win their first prize if they stayed still over both gradCPT functional runs, their second if they stayed still over the movie runs, and their third if they stayed still over the rest scans. A “low motion scan” was determined based on watching the participant during the scan through the eye-tracking camera and also by determining if the subject moved between frames by manual inspection by the MRI technologist and research staff. We note that we designed the prize system to be flexible across participants. For example, if it was deemed that a younger subject would respond positively to prizes, we would increase the number of prizes they could win up to five. Alternatively, if an older subject was not interested in prizes, we did not force them to pick out prizes before the scan. In our experience, this flexibility is vital during the actual MRI session, and allowing for it can help increase the number of usable scans.

We also performed a brief practice on the day of the scan to reacquaint the participants with the scanner. This took place 15-20 minutes before the start of the MRI protocol and lasted 5 minutes. Scanner sounds were played while the participant lay in the mock scanner. Feedback was given as appropriate regarding head movement by research staff (MoTrak was not used).

Finally, we note that two of the subjects in the informal mock scan group (subjects 6 and 7) used the weighted blanket and prize system in the MRI session. Because these subjects did not receive a formal mock scan, we consider them to be a part of the informal mock scan group. Two of the subjects in the formal mock scan group (subjects 8 and 9) used an early version of a formalized mock scan protocol (and also used the weighted blanket and prize system in the MRI session). This early version of the mock scan used the same equipment, software, hardware, and the same general steps, but was approximately 5-10 minutes longer. Specifically, these subjects completed the mock scan protocol without picking a prize (i.e. they did not complete step four in Table 2), and the mock scans in steps 3, 5, and 6 were conducted for five-minutes.

### MRI session

After a localizer, participants underwent the following scans (acquisition parameters described in detail below; note that we also describe the task-based scans in more detail in a different section below): anatomical magnetization prepared rapid gradient echo (MPRAGE); T1 fast low angle shot (T1 FLASH); two runs of an attention task (the gradual onset continuous performance task (gradCPT); [41–43]); T2-weighted 3D fast spin echo image (T2 SPACE); four movie runs, in which participants were shown clips of an actress with and without eye-contact, with and without speech; and two resting-state runs (Table 3). The total length of scan time in the protocol was 61 minutes, 13 secs. Note that eye-tracking data are being collected in this study; this typically adds approximately 5-10 minutes for participant setup and 5-10 minutes for calibrating/validating the eye-tracker before data collection. Hence, typical scan times (including all setup, interactions with participants in between scans, etc.), ranges from 80 to 90 minutes.

**Table 3.** The scans conducted in this study. Note that the scans are listed in the order they are acquired. The scan times for all functional runs include shimming and magnet equilibration.

| Scan | Duration |
| --- | --- |
| 1. Localizer | 10 secs |
| 2. MPRAGE | 5 mins 45 secs |
| 3. T1-FLASH | 2 mins 39 secs |
| 4. gradCPT run 1 | 5 mins 26 secs |
| 5. gradCPT run 2 | 5 mins 26 secs |
| 6. T2-SPACE | 11 mins 7 secs |
| 7. Movie run 1 | 4 mins 57 secs |
| 8. Movie run 2 | 4 mins 57 secs |
| 9. Movie run 3 | 4 mins 57 secs |
| 10. Movie run 4 | 4 mins 57 secs |
| 11. Resting-state run 1 | 5 mins 26 secs |
| 12. Resting-state run 2 | 5 mins 26 secs |
| Total scan time | 61 mins 13 secs |
| Total scan time with eye-tracker setup, time between scans, etc. (approximate) | 80-90 mins |

Of note, all subjects were able to complete the entire protocol, except one participant in the formal mock scan group and one participant in the replication group. The participant in the formal mock scan group had mean FFD values < 0.10 mm for all functional runs, but did not want to complete the final rest scan and was unable to be redirected. The participant in the replication group wanted to come out of the scanner with approximately two minutes left in the final rest scan. The mean FFD of this scan was high (0.615 mm FFD). For the purposes of analyses requiring us to classify a scan as low or high motion, we considered both of these scans (from the formal mock scan group participant and the replication group participant) to be high motion scans. For the replication group subject, we used the mean FFD obtained from the partial scan when necessary in calculations.

In addition, for the informal mock scan, formal mock scan, and replication subjects, if a participant exhibited a gross head movement, was not complying with task instructions, etc., we repeated the appropriate scan (since the goal of ongoing study is to obtain as much high quality data as possible) if time permitted. For this study, we had to repeat a scan three times: one subject from the informal mock scan group (subject 6, movie 4; mean FFD of first scan = 0.3323 mm; mean FFD of repeat scan = 0.4433 mm; Table 4); one from the formal mock scan group (subject 18, movie 3; mean FFD of first scan = 0.2969 mm; mean FFD of repeat scan = 0.2167 mm); and one from the replication group (subject 4, movie 3; mean FFD of first scan = 0.2007 mm; mean FFD of repeat scan = 0.1985 mm). For each subject, we used the scan with lower mean FFD for the analyses described here.

**Table 4.** Mean FFD values of participants in which scans had to be repeated. Note that “subject number” refers to the number listed in Fig. 4 for the informal and formal mock scan groups and in Fig. 6 for the replication group, respectively.

| **Group of participant**<br>**(subject number)** | **Repeated scan** | **Original mean FFD** | **Repeated scan FFD** |
| --- | --- | --- | --- |
| Informal mock scan group<br>(subject 6) | Movie 4 | 0.3323 mm | 0.4433 mm |
| Formal mock scan group<br>(subject 18) | Movie 3 | 0.2969 mm | 0.2167 mm |
| Replication group<br>(subject 4) | Movie 3 | 0.2007 mm | 0.1985 mm |

**Fig. 4.**
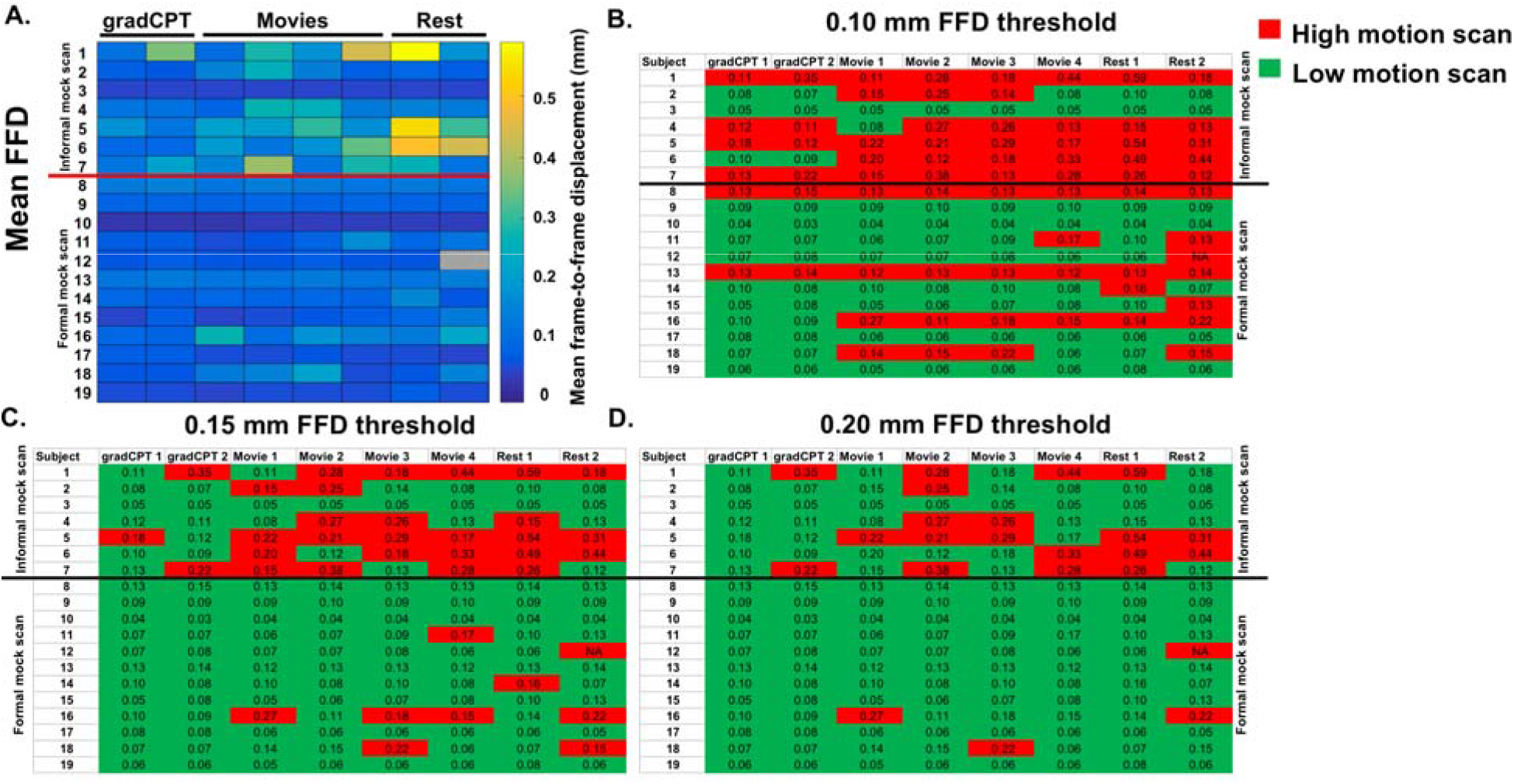
The formal mock scan group has fewer high-motion scans compared to the informal mock scan group. For all plots (a-d), each row represents a different participant; each column indicates a different scan condition (scan labels are shown at the top of each plot). Participants in the informal mock scan group are shown in the first 7 rows; those in the formal mock scan group are shown in rows 8-19. The line separating participant 7 and 8 divides participants into their respective group; group labels are shown off to the side of each plot. Note that the data are presented in functional run order: the gradCPT scans are the first functional scans conducted; the rest scans are the last functional runs conducted. Fig. 4a): A matrix of the mean FFD values for each participant. The color bar is scaled such that blue colors indicate a lower mean FFD for that scan; lighter colors (yellow) indicates a higher mean FFD for that scan. Fig. 4b–d): The same data from Fig. 4a are being plotted, except scans are classified as being either a “high motion” scan (i.e. the scan had a mean FFD above the threshold shown at the top of each plot) or a “low motion” scan (the scan had a mean FFD below the threshold). High motion scans are shown in red; low motion, in green. The mean FFD value for each scan is shown in each cell. Note that for visualization, we have rounded the mean FFD value to two significant figures; when we classified a scan as low or high motion, we used four significant figures. Also note that in all figures, subject 12 (in the formal mock scan group) did not complete the last rest scan. In Fig. 4a, this scan is shown with grey hatched lines; in Fig. 4b–d, we considered this a high motion scan and have colored it red (FFD = frame-to-frame displacement; mm = millimeters)

### Acquisition parameters

All subjects were scanned on a 3 T Siemens Prisma system at the Yale Magnetic Resonance Research center. We acquired a high-resolution T1-weighted 3D anatomical scan using a magnetization prepared rapid gradient echo (MPRAGE) sequence with the following image parameters: 208 contiguous slices acquired in the sagittal plane, repetition time (TR) = 2400 ms, echo time (TE) = 1.22 ms, flip angle = 8°, slice thickness = 1 mm, in-plane resolution = 1 mm × 1 mm, matrix size = 256 × 256. A T1-weighted 2D anatomical scan was acquired using a fast low angle shot (FLASH) sequence with the following image parameters: 75 contiguous slices acquired in the axial-oblique plane parallel to AC-PC line, TR = 440 ms, TE = 2.61 ms, flip angle = 70°, slice thickness = 2 mm, in-plane resolution = 0.9 mm × 0.9 mm, matrix size = 256 × 256. A T2-weighted 3D fast spin echo image was acquired using a sampling perfection with application optimized contrasts using different flip angle evolution (SPACE) sequence with the following image parameters: 208 slices per slab acquired in the saggital plane, TR = 3200 ms, TE = 316 ms, slice thickness = 1 mm, in-plane resolution = 1 mm × 1 mm, matrix size = 256 × 256 × 208.

Functional images were acquired using a multiband gradient echo-planar imaging (EPI) pulse sequence with the following image parameters: 75 contiguous slices acquired in the axial-oblique plane parallel to AC-PC line, TR = 1000 ms, TE 30 ms, voxel size = 2.0 mm^3^, flip angle = 55 degrees, slice thickness = 2 mm, bandwidth = 1894 Hz/pixel, matrix size = 110 × 110, field of view = 220 mm, multiband factor = 5.

### Description of functional runs

All tasks were presented using Psychtoolbox (version: 3.0.14; http://psychtoolbox.org/; MATLAB version R2018a) on a Lenovo IdeaPad 720S computer, with Ubuntu 16.04 LTS installed.

#### gradCPT

gradCPT is a continuous attention task that has been described in detail elsewhere [41–43]. Briefly, participants viewed grayscale images of city and mountain scenes presented at the center of the screen. In each trial, an image transitioned from one to the next through linear pixel-by-pixel interpolation. Each transition took 1000 ms. For 1000 ms the current scene transitioned from the previous scene, and for the next 1000 ms it transitioned to the next. Subjects were told to respond by pressing a button for city scenes and to withhold button presses for mountain scenes. City scenes occurred randomly 90% of the time. As in previous uses of gradCPT, accuracy was emphasized without reference to speed. All participants practiced outside of the scanner for 30 seconds to gain familiarity with the task.

#### Movies

The movie runs utilized a novel version of a free-viewing Selective Social Attention task [44, 45], in which an actress is situated at the center of the screen and is surrounded by four toys in corners of the screen (Fig. 3). Four conditions were used: The first clip was a direct gaze condition with speech, in which the actress spoke in full sentences (e.g. “Have you ever seen a monkey? Monkeys eat bananas, swing in trees, and chase each other.”) and used child- and adolescent-friendly language while smiling and making eye-contact with the camera. The second clip was a direct gaze condition with no speech, in which the actress smiled directly at the viewer while not speaking. The third clip consisted of the actress looking down at the table while speaking in full sentences while smiling (i.e. this is similar to the first clip except no eye-contact was made with the viewer). The fourth clip consisted of the actress looking down at the table while smiling and not speaking. Each clip lasted two minutes and was shown twice over four runs, such that eight clips were shown total. In between clips during each run, a white fixation cross on a black background was shown for 15 seconds. Clip order was counterbalanced across participants.

#### Resting-state

Subjects were instructed to keep their eyes open, relax, and think of nothing in particular while they viewed a white fixation cross on a black screen.

### Quantification of motion

To quantify subject motion, we calculated the mean frame-to-frame displacement (FFD) of the patient’s head across each functional run. We estimated a set of motion parameters using SPM8 [46] to obtain a transformation matrix *T* at frame *i* that allowed us to map the average Euclidean distance that the center of gravity (COG) of each frame moved. We then calculated the average movement over all frames acquired during a functional run. In equation form, we computed

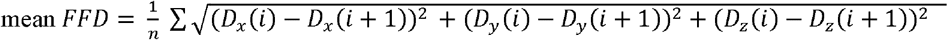

where *n* = number of frames, *i* = time point, and *D*_*x*_(*i*), *D*_*y*_(*i*), and *D*_*z*_(*i*) are the x, y, z elements of *T*(*i*) * COG at time *i*, respectively, leaving us with a scalar estimate of motion (mean FFD in mm) for each functional run.

We next determined the number of high and low motion scans in the informal mock scan and formal mock scan groups. If a participant’s scan had a mean FFD above a threshold, it was designated as a “high motion” scan; if it was below the threshold, it was considered a “low motion” scan. Mean FFD thresholds of 0.10, 0.15, and 0.20 mm were used, as thresholds of this magnitude have been shown to limit the impact of motion [25, 47] while allowing for sample sizes of adequate size in children/adolescents [47, 48] and in those with a disorder [10, 49, 50].

### Statistics

To determine if there were demographic differences among the informal mock scan and formal mock scan groups, we conducted two-sample *t*-tests; to determine if there were differences in the number of females per group, we used a two-tailed Fisher Exact Probability test.

In terms of the scanning data, we used a Chi-square test of association to determine if there was a significant difference in the number of high and low motion scans between the informal mock scan and formal mock scan groups. To determine if the mean FFD values differed due to scan condition (movies, gradCPT, and rest), we calculated Hedge’s *g* to determine effect sizes [51] (which is the preferred estimate of effect size when sample sizes are small [52] and unequal; [53]), and we used the criteria suggested by Sullivan and Feinn [54] for effect size interpretation. To determine statistical significance, we also performed two-sample *t*-tests on the average mean FFD value for a condition (i.e. for the rest condition, we averaged the mean FFD value from both rest scans for a participant). In addition, we conducted a two-sample *t*-test on the average mean FFD value for a participant across the entire session. Significance was assessed at a *P*-value < 0.05 after correcting for multiple comparisons using the Benjamini-Hochberg procedure [55].

We used a similar approach to that outlined above when examining the replication group. That is, we use a Chi-square test of association to determine if there was a significant difference in the number of high and low motion scans between the informal mock scan and replication groups, Hedge’s *g* to determine effect sizes, and performed a two-sample *t*-tests on the average mean FFD value for a condition, assessing significance at *P*-value < 0.05 after correcting for multiple comparisons using the Benjamini-Hochberg procedure.

## Results

### High and low motion scans in the informal mock scan and formal mock scan groups

We first set out to determine if there were differences in motion among the formal mock scan group and informal mock scan group. The demographic characteristics of these groups were not significantly different in terms of age, sex, and measures of IQ (Table 1).

To give a sense of the distribution of mean FFD values of these two groups, we show the mean FFD of each scan through multiple visualizations. First, we show the data in matrix format, in which each cell is colored according to the mean FFD of the participant for that scan (Fig. 4a). Visual inspection reveals most of the high mean FFD scans belong to participants in the informal mock scan group. Performing a similar analysis, except denoting each scan that is above a mean FFD threshold of 0.10 mm (Fig. 4b), 0.15 mm (Fig. 4c), and 0.20 mm (Fig. 4d) as a high motion scan, revealed that most of the high motion scans were from the informal mock scan group. For example, at a threshold of 0.10 mm, 71.4% of the scans from the informal mock scan group were high motion; only 32.3% of the scans from the formal mock scan group were high motion. The difference in high motion scans was statistically significant (Pearson Chi-square = 21.76, *P* < 0.001). We obtained similar results when using thresholds of 0.15 mm (50% of the scans from the informal mock scan group were high motion; only 9.38% of the scans from the formal mock scan group were high motion; Pearson Chi-square = 31.69, *P* < 0.001) and 0.20 mm (33.9% and 4.17% of the scans were high motion in the informal mock scan and the formal mock scan groups, respectively; Pearson Chi-square = 24.4, *P* < 0.001). Hence, imposing a movement threshold in this sample, a relatively common procedure to limit the impact of motion on functional connectivity analyses (e.g. [47, 48, 50]), revealed that the formal mock scan group had more low-motion data relative to the informal mock scan group.

### Comparison of mean FFD values between informal mock scan and formal mock scan groups

We next assessed if there were any differences in scan conditions between the informal mock scan and formal mock scan groups. Rather than classifying a scan as either high or low motion, we analyzed the mean FFD values for each participant (Fig. 5). We found that across conditions, the formal mock scan group tend to have lower mean FFD values relative to the informal mock scan group. Across all conditions, the size of the effect tended to be large (Table 5). The differences in group movement were significant across all conditions tested (except gradCPT; Table 6), such that the formal mock scan group had a statistically lower mean FFD group average compared to the informal mock scan group.

**Fig. 5.**
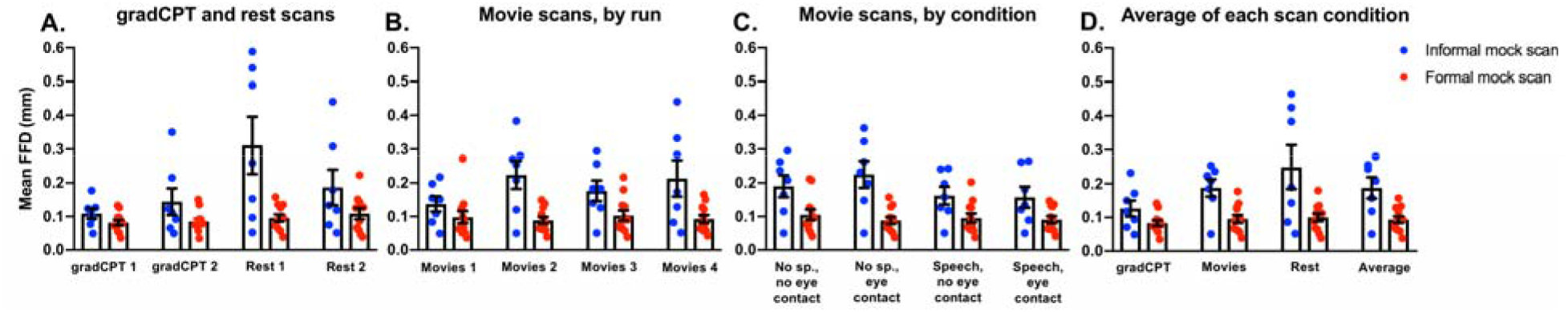
The formal mock scan group has a lower group average mean FFD compared to the informal mock scan group. For all plots (a-d), the scan condition is shown below the x-axis; the mean FFD (mm) is shown on the y-axis. Participants in the informal mock scan group are shown as blue circles; participants in the formal mock scan group as red circles. The average mean FFD for each group/condition is shown as a bar; error bars correspond to standard error of the mean. Fig. 5a): mean FFD values for gradCPT and rest scans. Fig. 5b): mean FFD values for the movie scans. Note that these are shown in order: movie 1 is the first functional run for the movie condition; movie 4 is the last functional run for the movie condition. The clips shown within each run were counterbalanced. Fig. 5c): plotting the same data as in Fig. 5b, except the mean FFD over each clip is shown. Of the four conditions listed under the x-axis, each is shown twice over the four runs for a total of eight clips. Fig. 5d): the average mean FFD value for each condition. For the movie runs, we averaged the data shown in Fig 5b. The “average” shown under the x-axis on the right of the plot represents the average mean FFD value over all eight functional scans (FFD = frame-to-frame displacement; mm = millimeters; sp. = speech)

**Table 5.** The effect size of the difference between the average mean FFD value of the formal mock scan and informal mock scan groups. The left-most column indicates the condition being compared. The right-most column indicates the qualitative description of the effect size based on the following scale: small (d⍰⍰≥⍰⍰0.2), medium (d⍰⍰≥⍰⍰0.5), and large (d ≥ 0.8) [54].

| Condition | Hedge's *g* | Effect size |
| --- | --- | --- |
| gradCPT 1 | 0.82 | Large |
| gradCPT 2 | 0.87 | Large |
| Movie run 1 | 0.63 | Medium |
| Movie run 2 | 1.89 | Large |
| Movie run 3 | 1.14 | Large |
| Movie run 4 | 1.33 | Large |
| Rest 1 | 1.58 | Large |
| Rest 2 | 0.81 | Large |
| Movies: No speech, no eye contact | 1.26 | Large |
| Movies: No speech, eye contact | 1.98 | Large |
| Movies: Speech, no eye contact | 1.18 | Large |
| Movies: Speech, eye contact | 1.16 | Large |
| Average gradCPT | 0.82 | Large |
| Average movie | 1.26 | Large |
| Average rest | 1.21 | Large |
| Average over all conditions | 1.08 | Large |

**Table 6.** Statistical significance of the difference between the average mean FFD value of the formal mock scan and informal mock scan groups.

| Condition | Degrees of | *t*-statistic | *P*-value (FDR corrected) | Formal mock scan |
| --- | --- | --- | --- | --- |

|  | **freedom** |  |  | **group mean FFD**<br>**significantly lower than**<br>**informal mock scan**<br>**group?** |
| --- | --- | --- | --- | --- |
| gradCPT | 17 | 2.05 | 0.0561 | No |
| Rest | 17 | 2.93 | 0.0094 | Yes |
| Movies | 17 | 3.7 | 0.0018 | Yes |
| Average across<br>all scans | 17 | 3.48 | 0.0029 | Yes |

### Results in replication group

Because the same individual (CH) led the mock scans and was responsible for communicating to the participants in between scans, it is possible that the results we obtained were specific to this individual and not due to the effects of the mock scan/prize system. To control for this potential confound, three other authors were trained in conducting the mock scans and administering the in-scanner prize system. Motion was then assessed in the subjects in the replication group who underwent mock-scanning by someone other than CH. Of note, we included 5 participants with ASD in the replication group, to assess if we were able to obtain low-motion data in this typically difficult-to-scan population. Demographic characteristics of this group were similar to the informal mock scan group in terms of age and IQ; after correction for multiple comparisons using the Benjamini-Hochberg procedure [55] there were no significant results (Table 1).

Similar to the formal mock scan group, we found we were able to obtain low motion scans in the replication group (Fig. 6a). At the 0.10 mm, 0.15 mm, and 0.20 mm FFD thresholds, we observed similar proportions of low motion to high motion scans as in the formal mock scan group (Fig. 6b–d). When we compared the mean FFD values obtained from a scan condition (i.e. gradCPT, movies, and rest), as well as the average across all scans within a participant, we again found that the replication group exhibited a statistically significant lower mean FFD compared to the informal mock scan group (except for the rest condition; Table 7). Calculation of effect sizes revealed that the difference in mean FFD relative to the informal mock scan group was large for all conditions, as well as the average of all scans (Table 7).

**Fig. 6.**
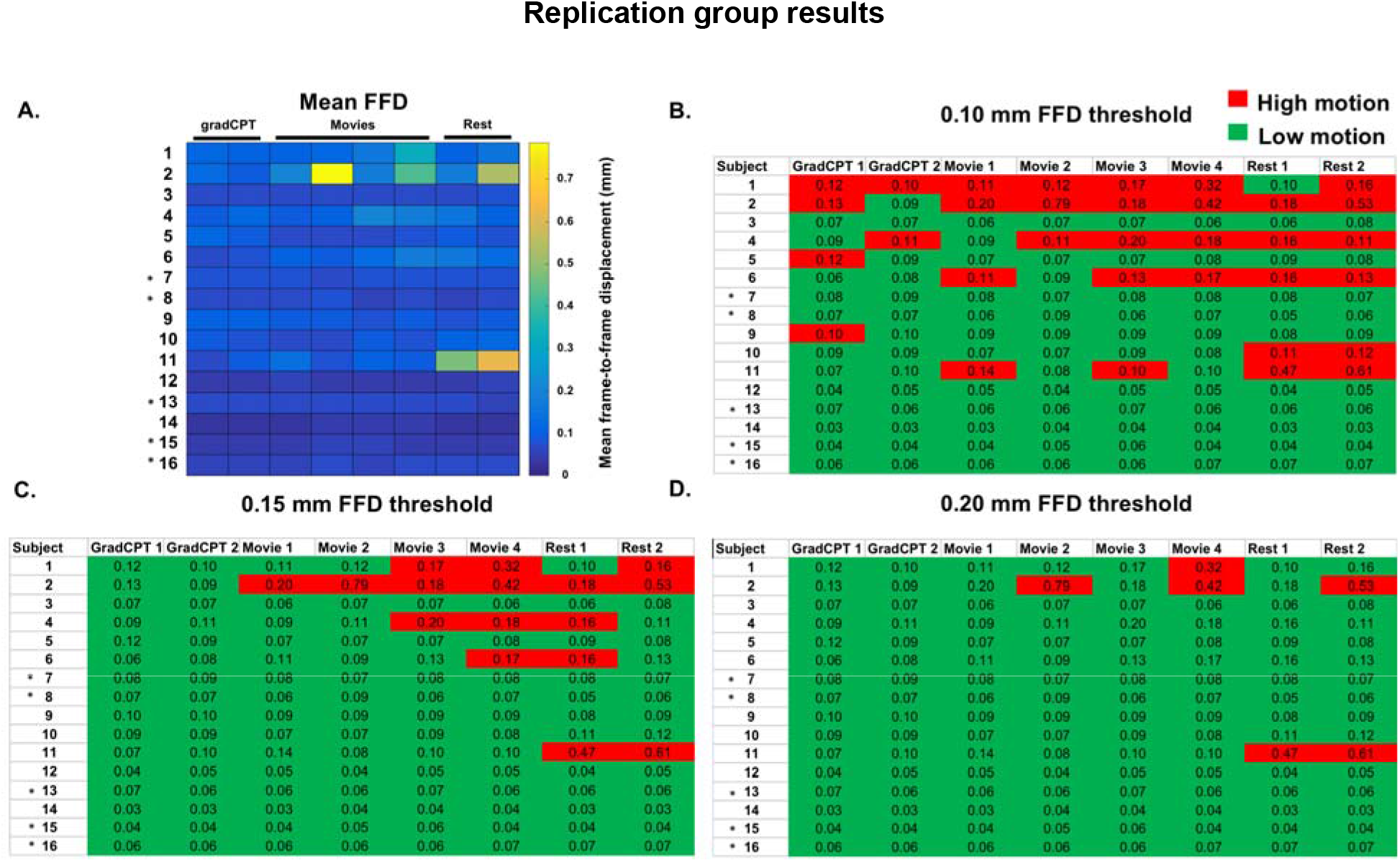
Low motion data can be obtained in a replication group of participants. For all plots (a-d), each row represents a different participant; each column indicates a different scan condition (scan labels are shown at the top of each plot). Participants with ASD are denoted by an asterisk (‘*’) to the left of subject number. Note that the data are presented in functional run order: the gradCPT scans are the first functional scans conducted; the rest scans are the last functional runs conducted. Fig. 6a): A matrix of the mean FFD values for each participant. The color bar is scaled such that blue colors indicate a lower mean FFD for that scan; lighter colors (yellow) indicates a higher mean FFD for that scan. Fig. 6b–d): The same data from Fig. 6a are being plotted, except scans are classified as being either a “high motion” scan (i.e. the scan had a mean FFD above the threshold shown at the top of each plot) or a “low motion” scan (the scan had a mean FFD below the threshold). High motion scans are shown in red; low motion, in green. The mean FFD value for each scan is shown in each cell. Note that for visualization, we have rounded the mean FFD value to two significant figures; when we classified a scan as low or high motion, we used four significant figures (FFD = frame-to-frame displacement; mm = millimeters)

**Table 7.**
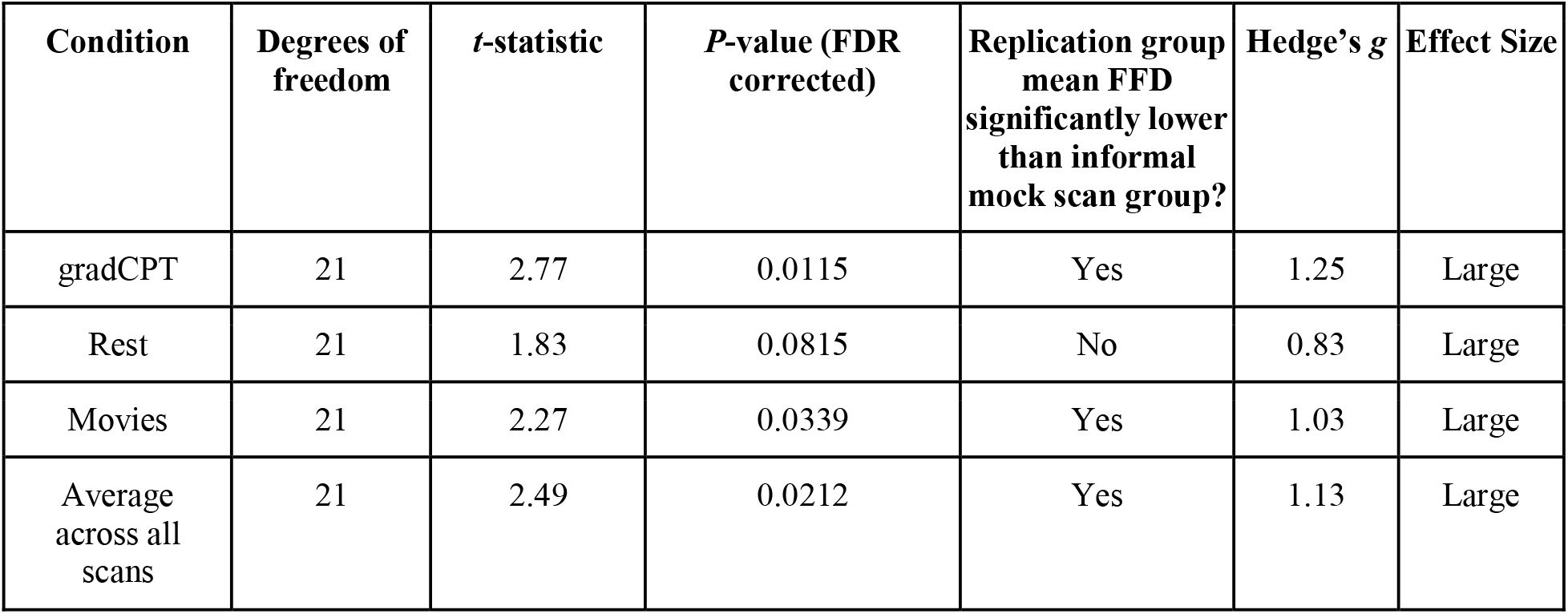
Effect size and significance testing of replication group compared to the informal mock scan group.

## Discussion

We have described steps our group has taken to limit in-scanner motion in pediatric participants undergoing an fMRI protocol. By implementing a mock scan protocol, utilizing an in-scanner incentive system that rewards participants for low-motion scans, and using a weighted blanket, we have significantly reduced movement in this sample. Crucially, we demonstrated that our formal mock scan group, with an intensive, systematic mock training session, exhibited lower motion data compared to a group of subjects who had only an informal mock scan experience. We determined these findings were robust to the experimenter conducting the mock and MRI scans. Taken together, these results indicate that our approach can be used to limit motion in pediatric participants undergoing a long MRI protocol.

### Comparison with other methods to reduce in-scanner motion

Scanning younger participants is a difficult task, especially those with a clinical condition or neurodevelopmental disorder. Given the effect of motion on functional connectivity analyses [14, 15], acquiring high-quality data can be challenging. As such, many groups have implemented steps before and during scans to limit participant movement. While these methods have proven useful, they are often not flexible enough to be applicable to all functional scans in a protocol (i.e. FIRMM can only be used in rest scans [29]), or they require researchers to provide feedback during a task-based scan, which might not be desirable for a given task [31]. In addition, using a method like *Inscapes* (i.e. a low demand clip of abstract shapes) can also help decrease motion, but it has been shown that this method affects connectivity and reflects some activation state [32]. The use of sedation or anesthesia to increase compliance has also been described [56–58], but it is well-acknowledged that such an approach can also affect connectivity [59–61]. The approach we have documented here circumvents these issues: it results in acquisitions with lower motion across both task and rest scans, does not require the use of additional stimuli during scans, and does not rely on sedation

### Adding to and extending the mock-scan protocol literature

The use of mock scanning protocols to increase MRI scan quality has been well-documented [33–36, 62]. Nevertheless, these studies have tended to use shorter MRI scan protocols (i.e. only 20-45 minutes) and have tended to acquire only structural and/or resting-state scans (or only a few task-based scans). We extend this work by showing the efficacy of a mock scan protocol in an MRI study that is longer (e.g. 60 minutes of total scan time) and uses a variety of task-based as well as rest scans, and we also compare the efficacy of our approach to an informal mock scan group from our same study, as well as a replication group of participants. To the best of our knowledge, these steps have not been taken before in papers documenting the use of mock scanning protocols to obtain low-motion functional data.

Directly comparing the efficacy of different mock scan protocols (and other procedures to limit motion) is difficult due to numerous differences among the actual MRI protocols in terms of number of scans, length of the MRI protocol, differences in the populations under study, different standards of what constitutes a “good scan,” etc. We therefore do so cautiously here, simply to place our results in the context of previous studies. In general, these previous studies have found that it is possible use a mock scan protocol and achieve low-motion data in approximately 70-80 percent of subjects (e.g. [33, 36]), while other groups prioritizing a scan of interest (i.e. a single structural image or a few resting-state runs) have reported success rates >85-90 percent (e.g. [37, 62]). Hence, our main contribution with this work is by showing that it is possible to achieve similar rates of low motion data, but in an MRI protocol that is longer and consists of many different types of scans. This is an encouraging finding, given that reliability of functional connectivity increases with more data per subject [24–28].

### Use with participants with ASD

We found that we were able to obtain low-motion data in five participants in the replication group who had ASD. Though the small number of subjects limits strong conclusions, these initial results are encouraging. Other groups have documented a number of methods to prepare participants with ASD for MRI scans, including mock scan protocols [37, 63, 64], as well as implementing motivational techniques and individualized prize systems (e.g., [37]). Our results add to this literature and provide additional tools that can be utilized to study this typically difficult-to-scan population. Using such tools is important, because we have known since the National Research Council report on autism [65] that some children with ASD are unresponsive to treatment, and the ability to include these children in neuroimaging studies may help us clarify differences in brain function that contribute to a lack of treatment response.

### Other motion considerations

We have described here one method to decrease motion. This work in no way invalidates, or precludes the use of, other approaches to decrease in-scanner movement. Indeed, depending on the goals of a specific study, one could use a mock-scan protocol before the MRI session. In the MRI session, a resource like FIRMM [29] could be used during the resting-state scans, specific feedback during or in between scans could be provided to participants [31], and/or resources like *Inscapes* [32] could be utilized as desired.

We also point out two other pertinent ways to decrease in-scanner motion that could be used in conjunction with a mock scan. The use of bite-bars is well known to many in the fMRI field and some labs use them in studies. Nevertheless, many participants describe bite bars as being extremely uncomfortable, and they have not been widely adapted by fMRI research groups. Custom head molds are another attractive option for reducing head-motion [66]. Nevertheless, it is unclear how younger participants, especially those with a disorder, would respond to such a head mold, as the work demonstrating the efficacy of this approach was conducted in healthy subjects that were older than the majority of subjects tested here. Given the influence of motion on connectivity measures, it is possible that some combination of these steps, along with post-hoc methods to detect and rid fMRI data of motion artifact, could be used to acquire high-quality functional connectivity data in populations that have been traditionally hard to scan.

Finally, we note the potential of our mock scanning protocol to act as a screening mechanism. Two participants were unable to complete the mock scan due to discomfort and a desire to end the session. They were unable to be directed, the mock scan session was aborted early, and the participants expressed that they did not want to complete the MRI protocol. Given the length of our protocol and the cost of scanning, excluding these two participants helped save both time and money. Hence, other groups could implement a similar mock scan procedure to help identify which subjects will give high motion and low quality data.

### Limitations and future directions

The small sample size of all three groups compared here is a primary limitation of this work. However, given that we achieved low motion data in two separate groups (i.e. the formal mock scan and replication groups), this increases confidence in the efficacy of the mock scan protocol. Another limitation is that because the sample described here is part of a larger, ongoing study, the staff conducting the mock scans, as well as the actual MRI scans, were not blinded, and subject assignment was not random. (Indeed, the pre- and in-scan steps we describe here were implemented in response to high-motion, low-quality functional scans.) Blinded, randomized, systematic studies could be conducted with larger samples to determine the effect of specific steps to limit motion, though this might be impractical—an investigator would have little incentive to sacrifice data quality by not utilizing an intervention (to achieve a true “control group”) to decrease motion if it is readily available. Lastly, having participants complete a formalized mock scan prior to their actual scan might not be feasible for all research studies due to scheduling and cost purposes. Nevertheless, future work could determine the efficacy of a more streamlined mock scan protocol (i.e. one that does not last as long as the one described here, one that does not rely on hardware and software monitoring participant head movement, etc.).

## Conclusion

In summary, we have described the implementation of a pre- and in-scan protocol to limit in-scanner motion of pediatric participants undergoing a long fMRI protocol. Used in conjunction with other methods, such a protocol assists in obtaining high-quality fMRI data in difficult-to-scan populations.

## Acknowledgements

The authors thank Abigail Greene for helpful comments on an early draft of this manuscript, Hedwig Sarofin and Cheryl McMurray for technical assistance during the MRI scans, Jitendra Bhawnani for technical assistance with task hardware and software, and Dave O’Connor for helpful discussions related to this work.

